# PanScreen: A Comprehensive Approach to Off-Target Liability Assessment

**DOI:** 10.1101/2023.11.16.567496

**Authors:** Manuel S. Sellner, Friedrich L. Joos, Alex Odermatt, Markus A. Lill, Martin Smieško

**Author notes:** Contributing authors.

## Abstract

Drug development projects are getting increasingly more expensive while their success rate is stagnating. Safety issues attributed to off-target binding represent a major reason for the failure of new drugs. Besides desired on-target binding, small molecules may interact with off-targets, triggering adverse effects. Therefore, the development of resource-efficient and cost-effective novel methods for early recognition of such issues becomes vital. Here, we introduce PanScreen, an online platform for the automated assessment of off-target liabilities. PanScreen combines structure-based modeling techniques with state-of-the-art deep learning methods to not only predict accurate binding affinities but also give insight into potential modes of action. We show that the predictions are approaching experimental accuracy found in public datasets and that the same technology can also be used for other research areas, such as drug repurposing. Such fast and inexpensive methods allow researchers to test not only drug candidates, but all small molecules that might come into contact with a human organism for potential safety concerns very early in the development process. PanScreen is publicly available at www.panscreen.ch.

## 1 Introduction

Chemicals are omnipresent in the environment due to the use of drugs, pesticides, fertilizers, combustion engines, waste water, and industrial by-products. Humans are in constant contact with their environment, leading to unevitable exposure to a wide variety of chemicals. The most common sources of human exposure to chemicals include food and water, air, personal care and pharmaceutical products, clothing, and household products [1]. Environmental exposure to chemicals has long been known to have adverse effects on humans, such as various cancers, infertility, other reproductive disorders, respiratory diseases, and allergic reactions [2–4]. Pharmaceutical products are specifically made to be ingested, injected, inhaled, or topically applied by humans. Thus, it is especially important to ensure their safety by recognizing and avoiding toxic effects.

Most small molecule drugs are designed to interact with one or more proteins in the human body by modulating their physiological behavior [5–7]. Sometimes, modulating the target protein inevitably leads to adverse effects e.g. by the disruption of essential cellular pathways. This effect is known as on-target toxicity [8, 9]. Often, however, drugs not only interact with their intended target protein, but also with other so-called off-targets, leading to possible side effects [7]. Such toxicities are estimated to account for up to a third of the attrition of drugs [10, 11]. In some exceptional cases, off-target binding can even be beneficial [12].

Investigation and identification of potential off-target toxicities is therefore highly important. This is true not only for the pharmaceutical industry but also for environmental chemicals that may end up in the human body. Experimental off-target profiling is usually expensive, slow, and labour- and resource-intensive [13–17]. On the other hand, computational methods are cheap and fast. The immense increase of available computing power during the past decade combined with the continuous improvement of computational methods has enabled in-silico tools to become a valuable addition to experimental testing. It is therefore not surprising that several tools aiming to predict off-target toxicities have been developed in recent years [18–23].

In-silico methods are used to predict not only off-target toxicities but also various endpoints. Some tools predict assay outcomes such as mutagenicity or skin sensitization [24–27], others predict clinical outcome [28, 29]. In off-target liability prediction, the underlying mechanism is usually based on undesired interactions between a small molecule and an off-target. This falls within the scope of drug-target interaction prediction [30–33]. In a toxicology setting, drug–off-target interactions usually represent molecular initiating events in an adverse outcome pathway [34]. However, drug-target interaction prediction is not restricted to toxicology and can also be used in drug development.

### 1.1 Ligand-based methods

One of the principles frequently applied in drug discovery, as well as predictive toxicology, is that chemically similar molecules exhibit similar properties. Similarity can thereby be defined in various ways, such as 2D similarity or 3D shape overlap [35, 36].

The advantage of ligand-based methods is that they only need a seed (or template) ligand structure as input. This allows them to be used in most drug development projects with at least one known initial hit. Additionally, ligand-based methods such as similarity searches are usually computationally inexpensive, leading to fast results. For these reasons, it is no surprise that ligand-based methods are routinely used in off-target prediction and drug development in general [37–40].

Although these methods work well in many cases, they have some inherent disadvantages. Two-dimensional approaches may be limited to a particular molecular scaffold, which can lead to a similarity search missing out on hits with a different structure. Furthermore, two molecules can be highly similar in 2D structure and/or 3D shape but still exhibit completely different activities to a given target. This phenomenon is known as an activity cliff [41, 42]. In such cases, more detailed analyses are necessary to accurately assess the potency of a molecule.

### 1.2 Structure-based methods

In contrast to ligand-based methods, structure-based methods require the 3D structure or the primary sequence of the protein to be known. Structural data, especially well-resolved experimentally determined complexes, allow the protein-ligand complementarity to be decoded in very fine detail, enabling the methods to overcome the drawbacks of ligand-based methods [43, 44]. However, because of the generally higher computational cost of structure-based methods, they tend to be much slower than ligand-based methods.

One of the most commonly used methods in structure-based drug development is molecular docking, in which a small molecule is placed into the binding site of a protein while optimizing the molecular interactions between the two entities [45–48]. In docking, proteins are often treated as rigid bodies, while the ligands are allowed to be flexible. This has the disadvantage that induced fit effects cannot be captured and the result of the docking depends on the input conformation of the protein. While it is possible to allow the side chains (or even the backbone) of the binding site residues of the protein to be flexible, this introduces many more degrees of freedom, leading to an explosion of possible combinations and therefore computational cost.

One way of tackling this problem is the use of ensemble docking [49–51]. In ensemble docking, a ligand is docked to an ensemble of protein structures, usually coming either from molecular dynamics simulations or experimentally determined (X-ray, cryo-EM, or NMR) structures. Usually, the ensemble is selected to represent different conformational states of the protein binding site. This approach implicitly accounts for the protein flexibility while minimizing computational cost.

Our group has previously developed VirtualToxLab, a platform accessible via a simple Java application for the automated assessment of the toxic potential of small molecules [22, 23]. Relying on the concepts of structure-based modeling, it features a portfolio of 16 well-known and comprehensively prepared off-targets, an induced-fit-enabled docking program, and optimized scoring functions for each target. VirtualToxLab has been extensively used by academia, regulatory agencies, and industry partners for its predictions, especially for CYP450 enzymes and nuclear receptors.

### 1.3 Machine learning-based methods

In recent years, machine learning has emerged as a major challenger to classical ligand-and structure-based methods [52–54]. Many machine learning models such as random forest, support vector machine, or naive bayes have been developed to predict drug-target interactions [55–57]. With increasing computational power, deep learning models have become more popular. With the right architecture, deep learning models have the ability to outperform classical machine learning models [58]. Thus, many deep learning-based models have recently emerged that aim to predict drug-target interactions [59–63]. A popular method to improve the performance of deep learning models is ensemble deep learning, in which an ensemble of models is trained with the goal of improving the generalizability of the combined ensemble [64, 65].

Although deep learning models have great potential to substantially improve the predictive power of in silico tools, they are not trivial to train. The construction of the data set and the processing of input features are imperative for a robust and unbiased model. The incorrect handling of data sets and the use of too large molecular input feature vectors contaminated with irrelevant information have been shown to lead to an overestimation of model performance [66–68]. Therefore, it is essential to thoroughly evaluate all components of the model training process to create a reliable and accurate model.

### 1.4 Our contribution

In this work, we introduce PanScreen, an online platform for the prediction of off-target liability that is publicly available. Similar to its predecessor, VirtualToxLab, PanScreen features a portfolio of off-targets. All implemented off-targets are thoroughly prepared and validated. PanScreen applies an ensemble docking approach using multiple docking programs and processes the output with deep learning models. This not only allows accurate predictions of off-target interactions, but also provides structural insight into the potential mechanism of action. The platform presents a user-friendly web interface that allows easy access for researchers with various degrees of experience with in silico structure-based modeling. It is available free of charge for academic and non-commercial use.

Although the number of implemented off-targets is still limited, highly standardized processes of preparing protein structures and training deep learning models greatly facilitate the addition of new off-targets. We anticipate a rapid growth of the off-target portfolio in the very near future.

## 2 Results and Discussion

PanScreen is available as an online service at www.panscreen.ch. The web application was developed using the Django web framework and is served using an nginx webserver [69]. At the time of publication of this article, the platform contained 14 implemented off-targets (see Table 1 for a complete overview). Each implemented off-target consists of an ensemble of thoroughly curated protein structures (see Section 3.1 for more information).

**Table 1:** (Off-)targets currently implemented in PanScreen.

| Uniprot ID | Name | Family |
| --- | --- | --- |
| O60674 | Tyrosine-protein kinase JAK2 | Kinase |
| P03372 | Estrogen receptor alpha | Nuclear receptor |
| P04150 | Glucocorticoid receptor | Nuclear receptor |
| P07550 | Beta-2 adrenergic receptor | GPCR |
| P10275 | Androgen receptor | Nuclear receptor |
| P14416 | Dopamine receptor D2 | GPCR |
| P23458 | Tyrosine-protein kinase JAK1 | Kinase |
| P25103 | Substance-P receptor | GPCR |
| P28222 | 5HT receptor 1B | GPCR |
| P37231 | Peroxisome proliferator-activated receptor gamma | Nuclear receptor |
| P49286 | Melatonin receptor 1B | GPCR |
| Q08499 | Phosphodiesterase 4D | Hydrolase |
| Q92731 | Estrogen receptor beta | Nuclear receptor |
| Q9Y233 | Phosphodiesterase 10A | Hydrolase |

Users can upload query molecules in various, commonly used data formats (e.g., SDF, MOL2, or SMILES) and select the off-targets against which the submitted molecules should be screened. The uploaded molecule is converted to canonical SMILES using openbabel version 3.0.0 [70]. The canonical SMILES format allows for the unique encoding of molecular structures while conserving stereochemistry and protonation states. The canonical SMILES are then stored in a PostgreSQL database that connects the front end and the back end. If the same molecule has already been processed before, the results are fetched from the database and provided to the user without need of re-running the simulations. In case the submitted molecule has not been processed before, the back-end reads the canonical SMILES from the database and starts processing it.

First, the canonical SMILES is used to generate a 3D conformation of the molecule using Schrodinger’s LigPrep [71]. It is thereby up to the user whether the protonation states present in the input molecule should be preserved or whether protonation states at physiological pH should be automatically generated. The prepared molecule is then docked to the protein ensembles of the desired off-targets. Currently, PanScreen implements 3 different docking programs to generate poses and 2 programs to re-score and analyze the generated poses. Detailed information on this process can be found in Section 3.2. The information from molecular docking and the analysis of the generated poses are then fed into a consensus model (described in Section 3.3). For each implemented off-target, we developed a specialized consensus model. This was done to increase the performance of individual off-targets while avoiding the bias that could be introduced by providing the model with structural information of the protein [67]. Once all calculations are completed, the results are stored in the PostgreSQL database. Since the front-end is also connected to this database, the user will have immediate access to the results of the computations. An overview of the data flow in PanScreen is shown in Figure 1. The protein-ligand complexes generated by PanScreen can be viewed online and downloaded.

**Fig. 1:**
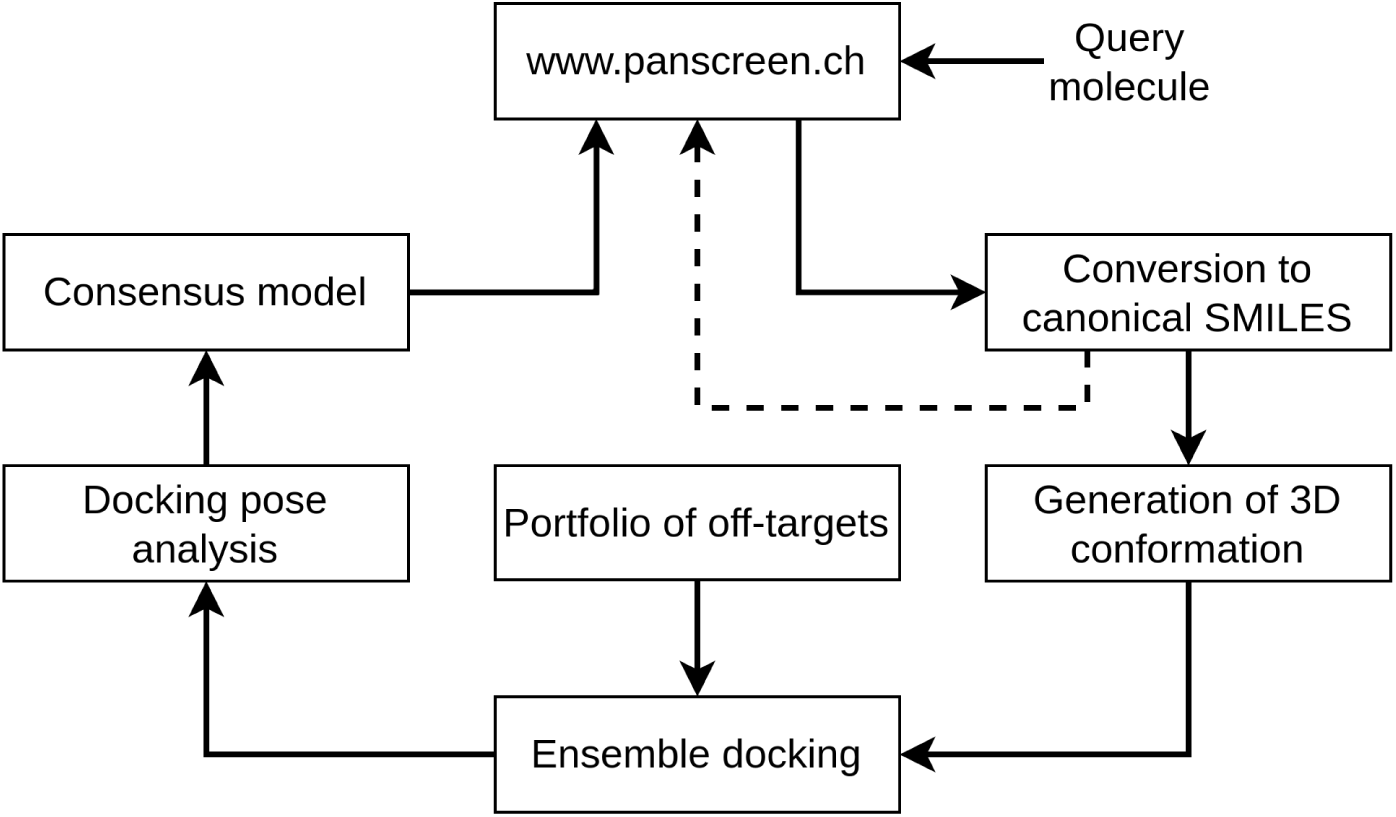
Flow of data in PanScreen. Once a compound is uploaded to PanScreen, it is converted to canonical SMILES and stored in a database. If the compound was processed before, its results are returned back to the user (dashed line). If the compound has not been processed before, it will be completely processed by the back end before the results are fed into the database and presented to the user.

### 2.1 Performance analysis

The validation performance of all implemented models can be found in Table 2 (the same analysis for smina, Glide, LeDock, and gnina can be found in the Supporting Information in Tables S2, S3, S4, and S5, respectively). The Pearson correlation coefficient (PCC) was above 0.70 for all models (except PPAR*γ*) with an average of 0.79. With exception of the melatonin receptor 1B and the estrogen receptor beta, all mean unsigned errors (MUE) were below 1.0 kcal/mol. The average MUE and root mean squared error (RMSE) were 0.83 kcal/mol and 1.12 kcal/mol, respectively. The area under the receiver operating characteristics curve (AUROC), calculated at an active/inactive threshold of 1.0 µM, was above 0.80 for all implemented off-targets with a mean of 0.89. This indicates very good performance for all models regardless of their protein family.

**Table 2:** Validation metrics of the implemented models. Shown are the Pearson correlation coefficient (PCC; higher is better), the mean unsigned error (MUE; lower is better), the root mean squared error (RMSE; lower is better), and the area under the receiver operating characteristics curve (AUROC; higher is better).

| Protein name | PCC | MUE<br>[kcal/mol] | RMSE<br>[kcal/mol] | AUROC |
| --- | --- | --- | --- | --- |
| Tyrosine-protein kinase JAK2 | 0.81 | 0.68 | 1.10 | 0.94 |
| Estrogen receptor alpha | 0.84 | 0.94 | 1.18 | 0.89 |
| Glucocorticoid receptor | 0.79 | 0.70 | 0.95 | 0.89 |
| Beta-2 adrenergic receptor | 0.79 | 0.92 | 1.17 | 0.91 |
| Androgen receptor | 0.81 | 0.79 | 1.03 | 0.88 |
| Dopamine receptor D2 | 0.75 | 0.66 | 0.87 | 0.88 |
| Tyrosine-protein kinase JAK1 | 0.81 | 0.55 | 0.85 | 0.94 |
| Substance-P receptor | 0.80 | 0.77 | 1.02 | 0.91 |
| 5HT receptor 1B | 0.77 | 0.83 | 1.06 | 0.85 |
| PPARγ | 0.68 | 0.89 | 1.33 | 0.84 |
| Melatonin receptor 1B | 0.72 | 1.06 | 1.35 | 0.84 |
| Phosphodiesterase 4D | 0.80 | 0.95 | 1.36 | 0.84 |
| Estrogen receptor beta | 0.75 | 1.00 | 1.29 | 0.84 |
| Phosphodiesterase 10A | 0.80 | 0.88 | 1.18 | 0.93 |
| Mean | 0.79 ± 0.05 | 0.83 ± 0.15 | 1.12 ± 0.17 | 0.88 ± 0.04 |

Plotting the predicted binding affinities against the experimentally determined affinities over all implemented off-targets (a total of more than 2800 predictions) revealed a PCC of 0.83 (see Figure 2). Moreover, more than 83% of the predictions were within 1 log unit of the true affinity. 15% of the predictions were between 1 and 2 log units from the experimentally determined binding affinity and only around 2% had a deviation of more than 2 log units.

**Fig. 2:**
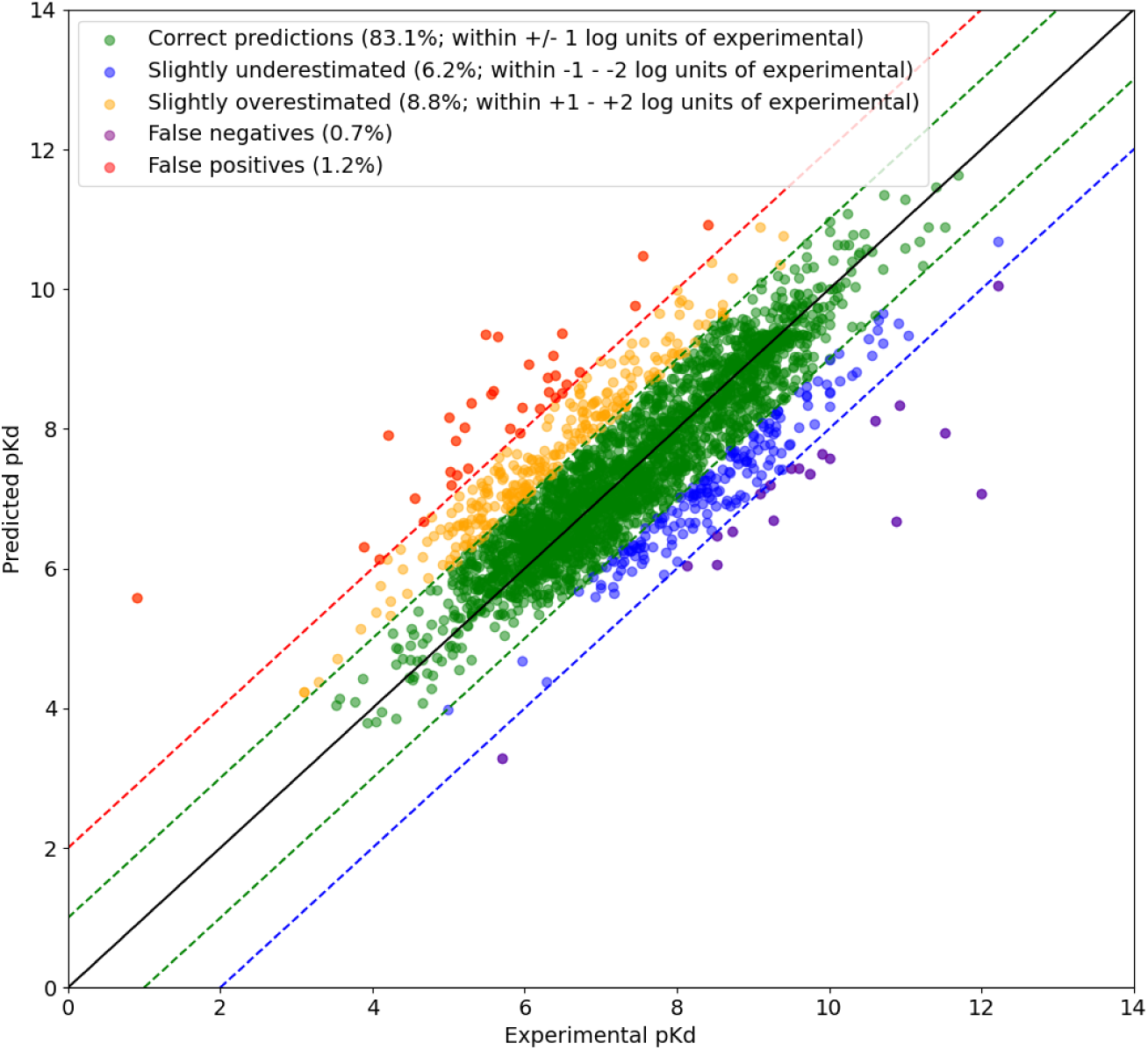
Correlation between predicted and experimental pKd values. The solid black line represents a perfect correlation. The green dashed lines represent a +/− 1 log unit deviation from the experimentally determined pKd values, the red and blue dashed lines represent a 2 log unit deviation from the experimental pKd values.

To investigate the meaning of these numbers, we analyzed the experimental accuracy in the datasets used to train the models. For this, we investigated all the data points with measured K_i_ and K_d_ values for every off-target currently implemented in PanScreen. We filtered the data for compounds that have been tested at least twice (around 2600 individual compounds) and calculated the maximum spread between the individual measurements. We found that 2083 compounds (80%) had a spread of less than 1 log unit, 375 (14%) were spread between 1 and 2 log units and 135 (5%) had a spread of more than 2 log units. These findings align very well with the accuracy of our predictions. In fact, when considering only compounds that have been measured at least 4 times (a total of 305 different compounds), we found a median and mean spread of 1.2 and 2.7 log units, respectively. This shows that our predictions reach the accuracy found in publicly available experimental datasets.

Regarding the previously discussed shortcomings of the ligand-based methods, we performed an in-depth investigation of how the structure-based approach implemented in PanScreen copes with matched molecular pairs (MMPs) [72–75]. MMPs are pairs of highly similar molecules that differ in only a few atoms. In some cases, MMPs have very different activities (binding affinities) despite their high degree of structural similarity. One such example is shown in Figure 3 where the molecule in subfigure a) is a very potent inhibitor of the Janus kinase 2 whereas the molecule in subfigure b) inhibits the Janus kinase 2 only very weakly. This is a prime example of an activity cliff caused by the removal of the hydroxyl group in Figure 3 b). Importantly, these compounds were not used in the training set of the respective model. For this example, there is an experimentally determined ΔΔG of 4.26 kcal/mol. The predicted binding free energies of Panscreen were within 1.0 kcal/mol of the experimentally determined values for both molecules and the predicted ΔΔG was 3.82 kcal/mol.

**Fig. 3:**
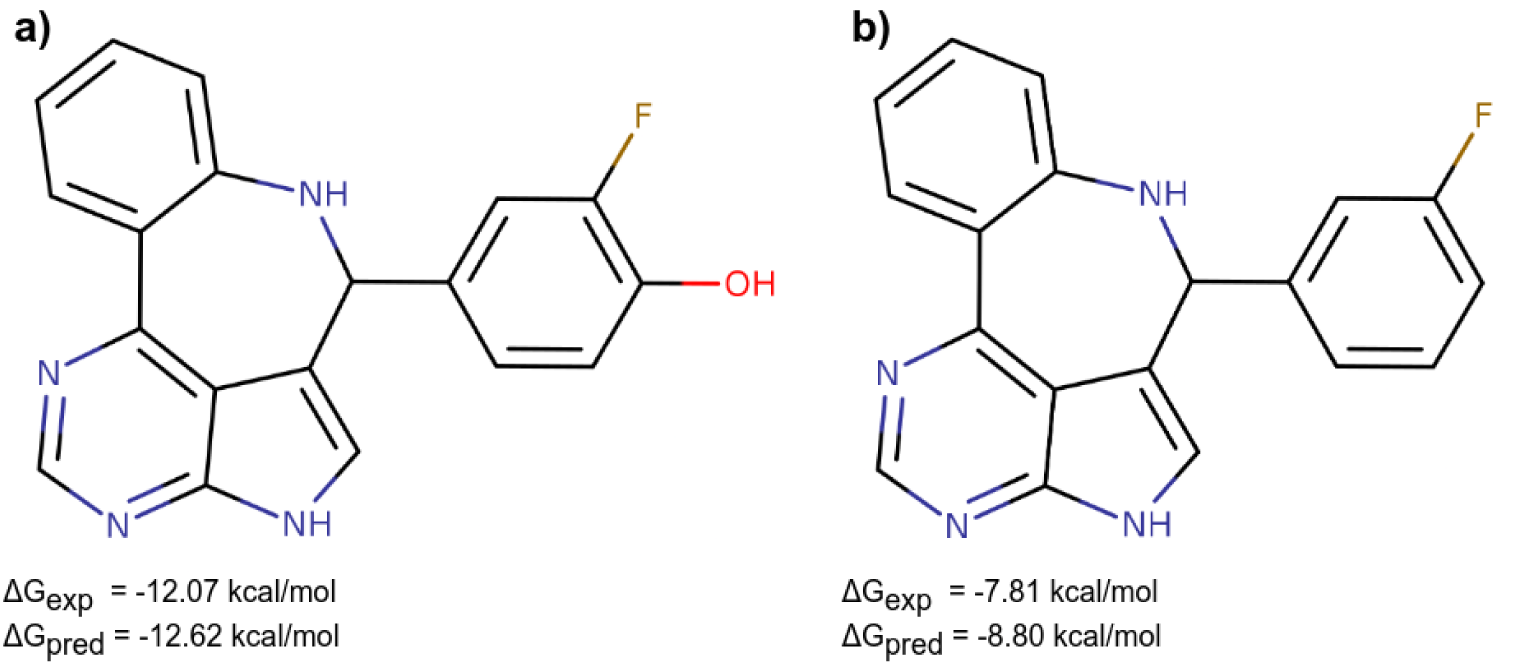
Example for activity cliff. a) Potent inhibitor of the Janus kinase 2, b) Weak inhibitor of the Janus kinase 2.

Figure 4 depicts a comprehensive analysis of all MMPs found in the validation sets of the models for all implemented off-targets. Here, we defined MMPs as molecules with a Tanimoto similarity of at least 0.7. A confusion matrix containing the results can be found in Table S1. Of the 3466 identified MMPs that were not part of the training sets, 2926 (84.4%) had a predicted ΔΔG within ±1.0 kcal of the experimental ΔΔG. Only very few MMPs (38; 1.1%) were overestimated by the models. However, a total of 502 MMPs (14.5%) were underestimated, whereof 110 (3.2%) had a predicted ΔΔG that was more than 2 kcal/mol lower than the experimentally determined one. In these cases, our models were not able to correctly predict the activity cliffs. Our analyses showed that in most of these MMPs, the docking programs were unable to correctly account for the key structural difference and thus distinguish between the two molecules. The docking scores for these MMPs usually had a ΔΔG of less than 1 kcal/mol and our models were not able to correct the predictions. Thus, these shortcomings are mainly due to limitations of the implemented docking programs.

**Fig. 4:**
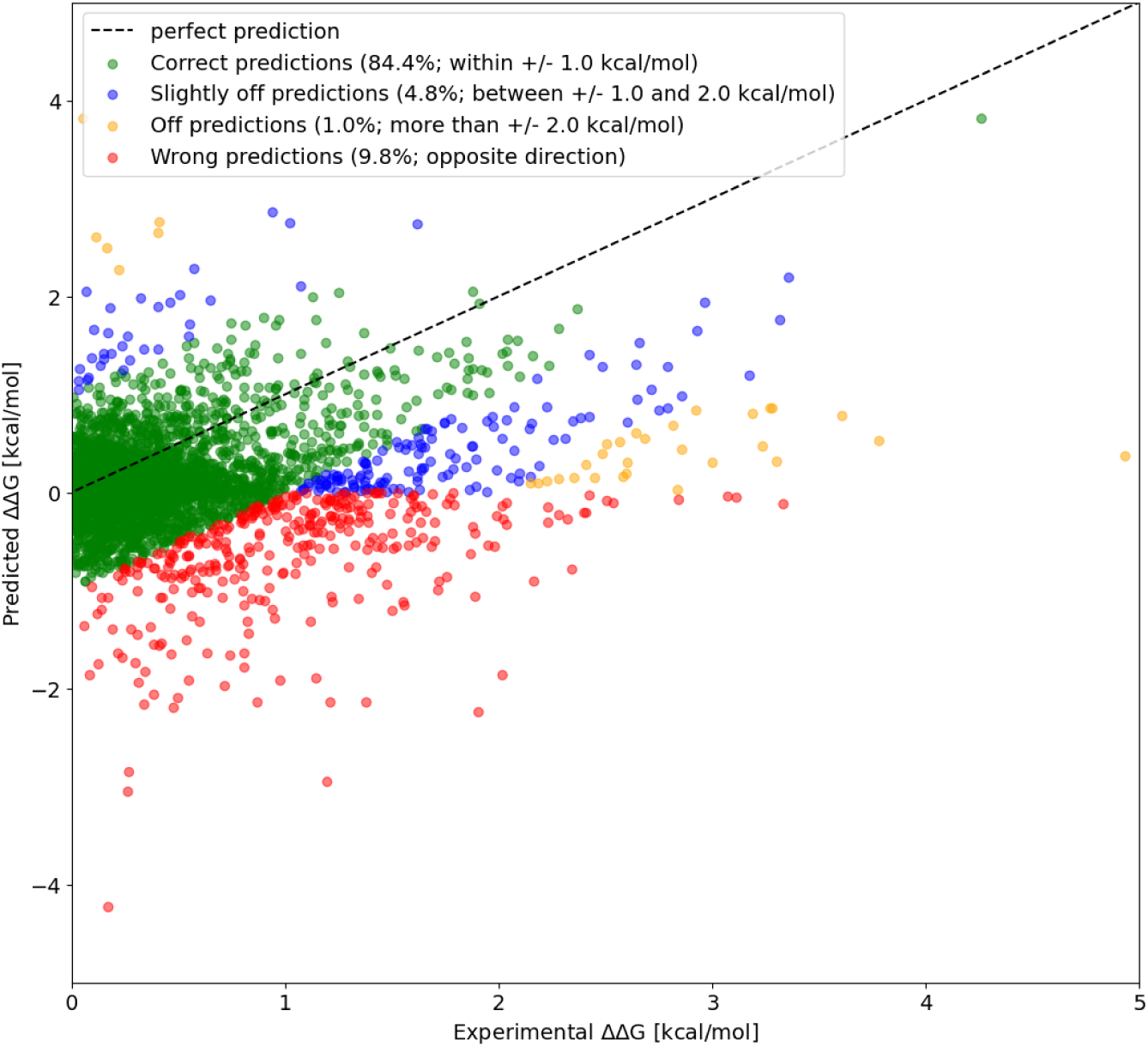
Analysis of matched molecular pairs. The black dashed line represents a perfect correlation between predicted and experimental ΔΔG values.

### 2.2 Screening performance

In order to further evaluate the quality of the predictions of our models, we screened the compounds contained in the Drugbank (after excluding ions and fragments) against the estrogen receptor alpha using our workflow [76]. We paid extra attention to only include compounds that were neither in the training nor in the validation set of the model. We filtered out all hits with an applicability score below 0.2 or with raised warning flags according to Section 3.3. The top 10 hits are shown in Table 3. The results showed that 5 of the top 10 hits have literature-confirmed activity on the estrogen receptor alpha. These compounds include steroids as well as non-steroidal structures. For the remaining 5 compounds of the top 10 hits, no reference for the activity at the estrogen receptor alpha could be found in the literature. We therefore analyzed the binding modes of these compounds and found that most of them could form reasonable interactions with the receptor, but most importantly had a good complementarity with the binding cavity (see Figures S1-S5 for the binding modes). This complementarity has been found to be an important factor in determining the quality of a binding pose [77]. Thus, there is a good chance that these compounds indeed bind to the estrogen receptor alpha. We therefore tested 4 of the remaining 5 predictions in a cell-based estrogen receptor alpha assay (DB08309 could not be obtained; see Section 3.5 for details on the assay). Detailed plots can be found in the SI in Figures S7-S9.

**Table 3:** Top 10 hits found by screening the Drugbank compounds against the estrogen receptor alpha. Only compounds that were neither in the training nor in the validation setused to train the model are shown.

| Compound name | $\Delta G_{\text{pred}}$ [kcal/mol] | Confirmed ERα activity |
| --- | --- | --- |
| DB06249 | -13.45 | Yes |
| DB08309 | -12.75 | n.a. |
| DB16139 | -12.52 | n.a. |
| DB01524 | -12.05 | Yes |
| DB02187 | -12.00 | Yes |
| DB00345 | -11.97 | n.a. |
| DB13866 | -11.96 | Yes |
| DB03882 | -11.96 | Yes |
| DB13591 | -11.74 | n.a. |
| DB15449 | -11.74 | n.a. |

One of the compounds, dihydrexidine (DB16139), is under investigation for the treatment of schizophrenia by acting as a dopamine D1 agonist [78]. It has been shown that estrogen receptor modulators such as raloxifene also have some activity on dopamine receptors [79, 80]. This indicates that there may be the possibility that compounds designed to bind to the dopamine receptor also interact with the estrogen receptor. Indeed, our study showed that dihydrexidine acts as a weak estrogen receptor alpha antagonist at a concentration of 10 *µM* (Figure S8).

Metopimazine (DB13591) is an antiemetic with antidopaminergic effects at the D2 and D3 dopamine receptors. Similar to dihydrexidine, it was also found to be a weak antagonist of the estrogen receptor alpha at 10 *µM* concentration (Figure S8).

DB15449 (citarinostat) is a histone deacetylase inhibitor. It consists of a triphenylamine analog, a linker, and a zinc binding group (Figure S6). The triphenylamine analog has a shape similar to that of cyclofenil analogues and therefore may also possess the ability to bind to the estrogen receptor alpha. If this was the case, citarinostat could act as a histone deacetylase inhibitor and estrogen receptor modulator hybrid [81–83]. Our experiments showed that citarinostat itself acts as a weak estrogen receptor alpha agonist at a concentration of 10 *µM* (Figure S9). However, it considerably enhanced the stimulation of estrogen receptor by 0.1 nM estradiol (Figures S7, S8), suggesting that this effect was mediated indirectly by histone deacetylase-related effects.

Lastly, for 4-Aminohippuric acid (DB00345), we could not find any estrogen receptor alpha activity in our assay (Figures S7-S9).

In total, the top 10 predicted hits were satisfactory with 5 literature-confirmed estrogen receptor alpha modulators, 2 compounds that could be confirmed to have weak antagonistic activity, and one compound that was shown to have weak modulatory activity, while one compound could not be experimentally validated. However, it needs to be noted that the transactivation assay to test for estrogenic activity was performed in intact cells, and it cannot be excluded that the compound did not reach the receptor due to efflux transporters. This shows that the tested model has a very good enrichment of the top N hits with confirmed or plausible molecules. Therefore, we believe that PanScreen could be advantageously applied for use cases other than off-target assessment, e.g. drug repurposing.

## 3 Methods

### 3.1 Selection and preparation of protein structures

All protein structures implemented in PanScreen have been experimentally determined and computationally prepared. We identified off-targets based on their Uniprot ID and used the associated crystal structures listed on Uniprot as the starting position [84]. The obtained crystal structures were then manually assessed using their entry in the PDB [85]. Only structures with co-crystallized ligands were considered while excluding fragments. We checked for mutations in the vicinity of the binding site and visually inspected the electron densities of the binding site residues and the co-crystallized ligands. All crystal structures with non-covalently bound co-crystallized ligands, an acceptable electron density at the binding site, and no mutations in the binding site were selected as potential ensemble candidates.

The goal of the ensemble selection was to minimize the size of the ensemble while maximizing the diversity of the contained structures. This was achieved by aligning all binding sites using the “align binding sites” routine that comes with Schrodinger Maestro version 2021-2 and selecting up to 4 structures with the highest binding site RMSD to each other [86]. The selected structures were then thoroughly prepared.

For the preparation of the protein structures, we used Schrodinger Maestro version 2021-2 [86]. We regenerated the crystal mates to ensure that there were no crystallization artifacts introduced by neighboring proteins in the crystal structure. In case there were binding site-remote mutations detected in the protein that could not affect the ligand binding mode, they were reverted to wild-type. We removed all crystallization adjuvants, but kept all physiological co-factors within a 12 Å radius around the ligand. The Protein Preparation Wizard within Maestro was used to assign bond orders, add explicit hydrogens, create zero-bond orders to metals, create disulfide bonds, convert selenomethionines to methionines, fill in missing side chains and loops, and generate protonation states at pH 7.4 ± 0.1 [87]. We then optimized the H-bond network at physiological pH and ran a minimization restrained to 0.3 Å. Finally, the structures were visually checked for any problems and fixed where necessary. A special focus was placed on the protonation states of aspartic acids, glutamic acids, and histidines, as well as flips of histidines, asparagines and glutamines.

It is well known that water can significantly influence the strength of a ligand binding to a protein [88–91]. Therefore, we modeled the binding site of the ensemble candidates in several different solvation states, depending on the availability of co-crystallized waters. When no co-crystallized water molecules were resolved, no solvation states were modeled. To select the final ensemble, we cross-docked all co-crsytallized ligands for a protein to the ensemble candidates in different solvation states. We calculated the lowest RMSD for each ligand-structure pair and selected the ensemble with the lowest average RMSD over all cross-docked ligands. It was therefore possible to get ensembles with more than one solvation state of a crystal structure, but we made sure that there were always at least 2 different crystal structures used in an ensemble. Figure 5 shows an overview of the complete ensemble generation process.

**Fig. 5:**
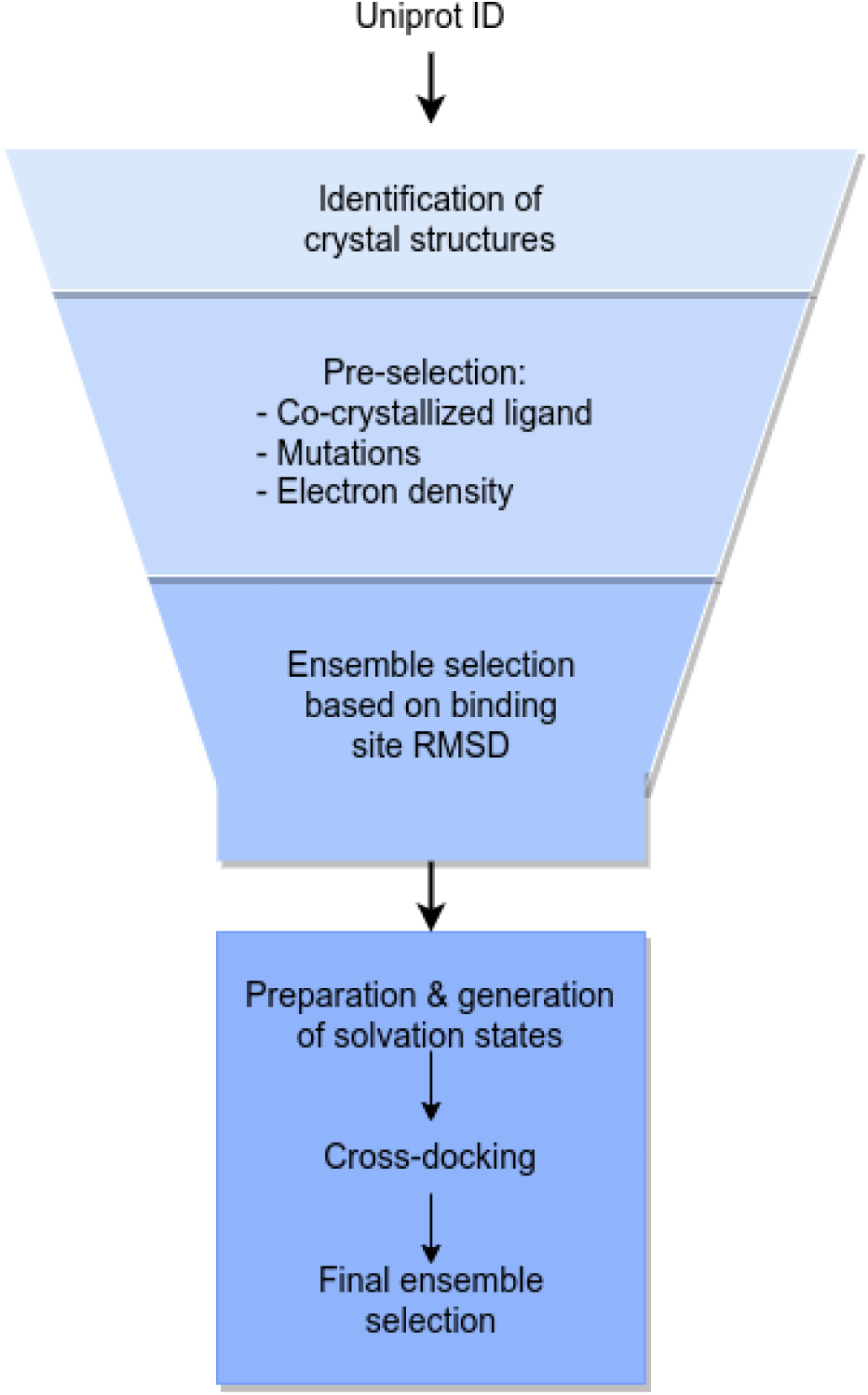
Process for the generation of ensembles. Crystal structures are selected based on a Uniprot ID and pre-filtered based on the electron density, co-crystallized ligands, and mutations. An initial crystal structure ensemble is generated by maximizing the binding site RMSD. The selected crystal structures are prepared in Schrodinger’s Maestro and different solvation states are modeled. Cross-docking to the solvation states is used to get the final ensemble.

### 3.2 Ensemble docking

PanScreen currently uses smina, Glide, and LeDock to generate docking poses and calculate accompanying scores [46, 47, 92]. For smina, the co-crystallized ligand was used to identify the binding site and the default buffer of 4 Å was added. We chose to generate up to 9 docking poses with an exhaustiveness of 16. For glide, we used single-precision docking to generate up to 10 docking poses. With LeDock, we generated up to 20 docking poses with a box constructed with a buffer of 6 Å around the co-crystallized ligand. Each ligand was docked to each structure in the ensemble using all 3 docking programs.

After generating protein-ligand complexes with the programs mentioned above, we used gnina to rescore all poses [93]. Gnina was run with the default model, the score only flag, with 2 CNN rotations, and an exhaustiveness of 16. Additionally, we used a model trained to predict binding affinities and generate protein-ligand interaction fingerprints based on po-sco. The training of this model followed the original publication [68]. This model was used to analyze all protein-ligand complexes generated by smina, Glide, and LeDock.

### 3.3 Consenus prediction

The calculated docking scores as well as the interaction fingerprints from the po-sco model were used to compute the final consensus prediction. This was done by training an individual consensus model for each implemented off-target. The docking scores of smina, Glide, LeDock, and gnina, as well as the affinity predicted by the po-sco model, were first converted to kcal/mol where necessary. We made sure that all affinities were less than or equal to zero by capping positive scores. In addition to the docking scores, we also calculated the standard deviation of the calculated docking scores over all generated poses for each program to estimate the uncertainty of the docking programs. The po-sco model also predicts an uncertainty estimation which was used for the same purpose. The docking scores and uncertainties were then passed through a radial basis function (RBF) expansion *r*(*x*) as shown in Equation 1, where *x* is the binding affinity predicted by a docking program or the po-sco model, the binning threshold set *c* is defined as *x_min_* = *c*_1_ *< c*_2_ *< < c_m_* = *x_max_* with *x_min_* = − 15 and *x_max_* = 0, and *m* is the number of bins. This number is subject to hyperparameter optimization and varies between models.

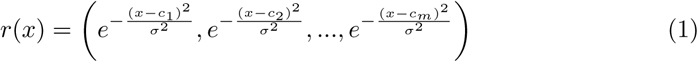

The definition of *σ* follows Equation 2 where *s* is also subject to hyperparameter optimization.

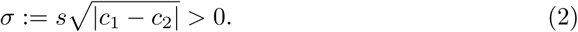

This means that we used in total 10 RBF representations as inputs which were all concatenated: 5 docking scores and 5 corresponding uncertainties. The interaction fingerprints for the best complexes from smina, glide, and LeDock were then concatenated with the expanded docking scores and uncertainties. Since it is not easily possible to objectively determine the “best” complex, we chose the one that had the best affinity predicted by the po-sco model. An overview of the input processing can be found in Figure 6A.

**Fig. 6:**
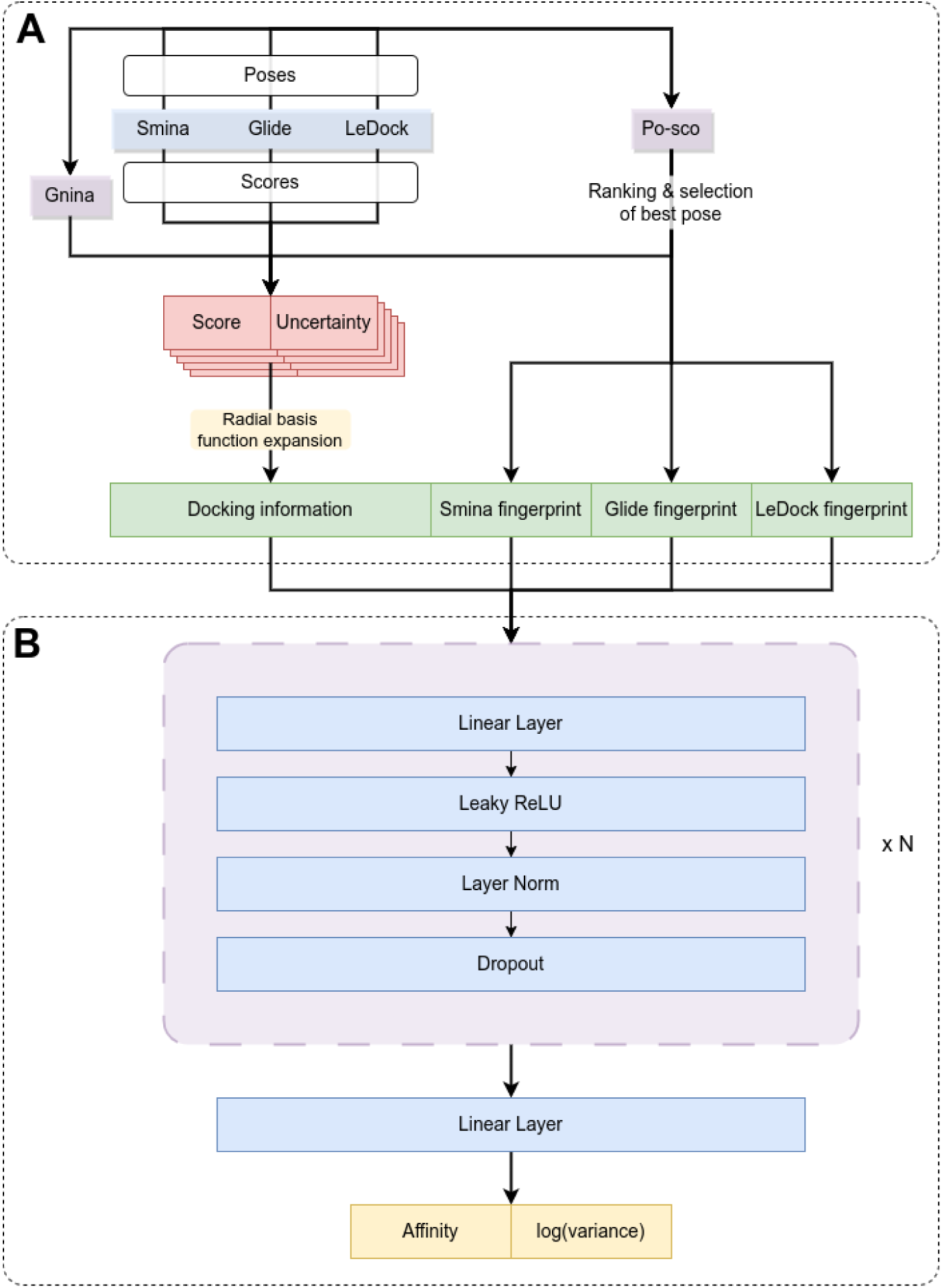
The full consensus model architecture. **A)** Processing of the inputs for the consensus model. Smina, Glide, and LeDock (blue) are used to generate docking poses and calculate docking scores and uncertainties (red). Gnina and the po-sco model (purple) are used to re-score the generated complexes, and their scores are combined with the ones of the docking programs (red). The scores and uncertainties are passed through a radial basis function expansion (yellow) to obtain the final docking information. The po-sco model is also used to generate interaction fingerprints for the best complexes generated by smina, Glide, and LeDock. The processed scores and the fingerprints are concatenated to form the final input of the model (green). **B)** Architecture of the consensus model itself. The processed input is passed *N* times through a linear layer followed by a leaky ReLU activation function, layer normalization, and a dropout node (purple). For the last layer, no activation function, layer normalization, or dropout is applied. The model predicts the affinity as well as the log variance.

The consensus model itself is a simple feed-forward neural network. A visual representation of its architecture can be found in Figure 6B. The processed inputs were passed through *N* feed-forward blocks. One block consisted of a linear layer, a leaky ReLU activation function, layer normalization, and a dropout node. The number of blocks (*N*) is subject to hyperparameter optimization and was in the range of 1 to 3. After the *N* feed-forward blocks, a single linear layer predicted the binding affinity and the log of the variance. The width of the linear layers was determined by a hyperparameter optimization for each target individually. During training, the predicted affinity and log variance were used to train the model using maximum likelihood estimation (minimization of the negative log likelihood loss). This is defined in Equation 3 where *θ* represents the model parameters, *n* is the number of samples in a batch, *y_i_* is the true label for sample *i*, *x_i_* is the input of sample *i*, and *p*(*·*) is the probability density function that gives the conditional probability of *y_i_* given *x_i_* and *θ*.

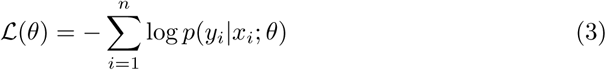

The neural network NN(·) predicts a distribution as 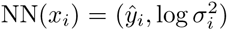 where *ŷ* is the predicted mean and log *σ*^2^ is the predicted log variance. The conditional probability *p*(*y_i_|x_i_*; *θ*) is then calculated as defined in Equation 4.

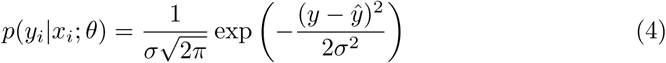

To increase the robustness of the model, we trained a total of 100 models with different seeds for the weight initialization for each off-target. The predictions of individual models were aggregated into a final prediction using a weighted mean based on compound similarities. This step was only done once all 100 models were trained. For each compound in the training set, we calculated the optimal weights for the 100 models using a multi-linear regression. For each unseen compound (from the test set or during inference), we calculated the Tanimoto similarities to all compounds in the training set based on Morgan fingperprints with a radius of 2 and size of 1024 bits. All training compounds with a similarity of *>* 0.75 to the unseen compound were selected as reference compounds. In case there were less than 5 training compounds with a similarity *>* 0.75 to the unseen compounds, the 5 training compounds with the highest similarity were chosen. The reference compounds and their similarities were then used to calculate a weighted average of the model weights according to Equation 5 where *w_j_* is a vector containing the model weights for the unseen compound *j*, *w_i_* are the optimal model weights for reference compound *i*, *N* is the number of reference compounds, and *s_ij_* is the similarity between reference compound *i* and unseen compound *j*.

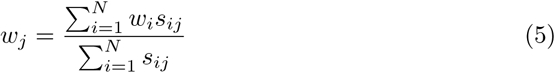

The data available for training and validation of the consensus models was usually limited, and the molecules did not cover the entire chemical space. Thus, estimating the applicability domain of the models is an essential part of the prediction. We did this by developing an applicability score. To calculate the applicability score, we first mapped all molecules used to train and validate a consensus model into 4-dimensional space using their smina, glide, LeDock, and gnina scores. We then constructed a convex hull around all data points in 4D space. When evaluating a query molecule, we mapped its docking scores into the same 4D space and checked whether it was within the hull. If it was, the applicability score was set to 1.0. If the query molecule fell outside the convex hull, we applied an exponential decay to the shortest distance between the molecule and the surface of the hull. The applicability score is therefore in the interval [0, 1] where higher scores indicate a better overlap with the applicability domain of the model.

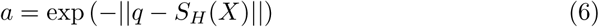

The calculation of the applicability score *a* is shown in Equation 6 where *q* is a query molecule, *X* represents all molecules used to train and validate the model, *S_H_* (·) is the surface of the convex hull around the set *X*, and ∥ · ∥ is the Euclidean norm.

In addition to the applicability score, we also introduced warning flags for our predictions. In total, there are 3 flags that could be raised: i) The compound is not binding (sum of smina, glide, and LeDock scores is *> −*8), ii) The compound could not be docked by all implemented docking programs, iii) The compound has unfavorable docking scores (sum of smina, glide, and LeDock scores is *> −*14).

### 3.4 Dataset generation

To train a model for a specific (off-)target, a dataset containing compounds with experimentally determined binding affinities are needed. For reproducibility, we developed a standardized routine to obtain and process these data. In the first step, the Uniprot ID of the target of interest is used to search for tested compounds on PubChem [94]. The obtained list is then filtered to only include data points with an activity type of either K_i_ or K_d_ and with an absolute affinity value (no values with indication “less than” or “greater than”). For compounds with multiple measurements, we first calculated the mean affinity and excluded all data points with measured affinities that deviate more than ± 30% from the mean affinity. Finally, we downloaded the 3D SDF files from PubChem for all remaining compounds. In some cases, there was no 3D structure available. This was mostly the case for very large and flexible compounds or for compounds with ambiguous stereochemistry. These compounds were excluded from the final dataset.

The final dataset was then sorted by decreasing affinity and every 5^th^ element was added to the validation set while the remaining elements were used for the training set. This approach was chosen to ensure a similar distribution of affinities between the training and validation set. Since the consensus model that was used to make the final predictions is agnostic of ligand structures, we did not pay attention to any structural similarities between the training and validation set. To deal with imbalances of high- and low-affinity compounds, we clustered all training compounds into 3 clusters with affinity thresholds of *<* 100 nM, *<* 1 *µ*M, and *>* 1 *µ*M. We then used a weighted random sampler to ensure the same numbers of high-, medium-, and low-affinity compounds per mini-batch.

### 3.5 Estrogen receptor alpha transactivation assay

#### 3.5.1 Seeding

24-well plates were coated with Poly L-Lysin solution (Sigma; 50 µg/mL in PBS) for 10 min and washed twice with PBS. HEK-293 cells were cultured in DMEM High Glucose (4,5 g/L) with L-Glutamine (BioConcept, Allschwil, Switzerland) with added 10% FBS, 1% penicillin-streptomycin, 1% HEPES-buffer and 1% 100x MEM non-essential amino acids. HEK-293 cells were seeded in poly-L-lysin-coated 24-well plates at a density of 100,000 cells/well in a volume of 500 µL medium/well and left to attach over night.

#### 3.5.2 Transfection

Plasmid DNA (390 ng/well ERalpha [95]; 25 ng/well CMVLacZ; 390 ng/well 4xERE Luc) was mixed with 37°C warm water to reach a volume of 30 µL/well. 10 µL/well of 37°C 2 M CaCl_2_ was added slowly. The mixture was pipetted dropwise into an equal volume of 37°C warm 2x BEST solution (8.0 g NaCl; 0.198 g Na_2_HPO_4_*·*7 H_2_O; 5.3 g N,N-Bis(2-hydroxyethyl)-2-aminoethanesulfonic acid, ad. 500 mL with H_2_O; pH = 7) and incubated at room temperature for 15 min for complex formation. Cells were transfected with 80 µL/well of the transfection solution. After 4 h the cell supernatant was removed and fresh medium (500 µL/well) was added. The cells were left to rest for another 20 h before the start of the treatment.

#### 3.5.3 Treatment

Treatment medium was cleared of estradiol by charcoal treatment: 50 mL medium were mixed with 0.2 g activated charcoal and shaken at 4°C for 1 h. The suspension was centrifuged at 4°C, 15000xg for 5 min and decanted into a new tube. This whole procedure was repeated 3 times for a total of 4 charcoal treatments. After the last charcoal treatment, the medium was sterile filtered with 0.2 µm Filtropur S 0.2 (Sarstedt, Nümbrecht, Germany). 1 h before the treatment, medium was changed to charcoal-treated medium to deplete the cells of estradiol. After 1 h the cell supernatant was changed to 500 µL charcoal treated medium containing the compounds of interest and 0.1% DMSO. To investigate antagonistic effects, after another hour 20 µL/well 2.6 nM estradiol was added for a total concentration of 0.1 nM estradiol in the well. After 24 h the cell supernatant was removed and cells were lysed for 5 min at room temperature with 60 µL Tropix lysis solution (Applied Biosystems, Foster City CA, USA). Cell lysate was kept at −80°C until further processing. Luciferase activity was determined by injecting 100 µL D-luciferin substrate solution [0.56 mM D-luciferin, 63 mM ATP, 0.27 mM CoA, 0.13 mM EDTA, 33.3 mM dithiothreitol, 8 mM MgSO_4_, 20 mM tricine (pH 7.8)] to 20 µL lysate prepared in pure grade white 96-well microtiter plates (BRAND, Wertheim, Germany) and measuring luminescence with the SpectraMax L Microplate Reader (Molecular Devices, San Jose, CA, USA). The respective *β*-galactosidase activity was measured with the same plates and device using the Galacto-Light Plus^TM^ kit (Applied Biosystems, Foster City CA, USA).

## 4 Conclusion

With the rise of increasingly accurate computational methods, in silico prediction of off-target interactions has become a viable tool to complement classical in vitro testing. In light of the FDA Modernization Act 2.0, we believe that it is the right time to further promote in silico methods due to their advantages in resource efficiency and cost effectiveness [96].

In this article, we present PanScreen, an online platform for the automated testing of off-target liabilities. At the time of writing, PanScreen features 14 (off-)targets of various protein families. Using a combination of structure-based modeling and artificial intelligence, all backed by profound knowledge in structural biology and medicinal chemistry, PanScreen is able to accurately predict binding affinities for diverse molecules. In addition to the predicted binding affinities, PanScreen offers possible binding modes as an explanation for the predictions. We also showed that PanScreen has the potential to detect activity cliffs between highly similar molecules. Due to the underlying technology, which is independent of a specific use case, our platform can be used not only for toxicology studies, but also for drug repurposing, selectivity assessment, and a wide range of other applications in the pharmaceutical and biomedical fields. To our knowledge, PanScreen is the first online platform that combines structure-based methods with deep learning to assess off-target interactions in a portfolio of highly curated proteins.

By providing PanScreen as a publicly available online platform, we hope to enable scientists of various backgrounds to use in silico off-target analysis with minimal effort and integrate the results in their own research.

## Supplementary information

This article has an accompanying supplementary file.

## Acknowledgments

This work was supported by the Biographics Laboratory 3R foundation and the Animalfree Research foundation. We gratefully acknowledge the support of NVIDIA Corporation with the donation of two RTX A5000 GPUs used for this research. This work was further supported (AO) by the Swiss Centre for Applied Human Toxicology Grant SCAHT-GL 25-01.

