## Supplementary Information for "PanScreen: A Comprehensive Approach to Off-Target Liability Assessment"

### Supporting Information: PanScreen: A Comprehensive Approach to Off-Target Liability Assessment

Table S1: Confusion matrix for the MMP analysis.

| | $\Delta\Delta G_{pred} < 1.0$ | $\Delta\Delta G_{pred} < 2.0$ | $\Delta\Delta G_{pred} > 2.0$ | **total** |
|---|---|---|---|---|
| $\Delta\Delta G_{exp} < 1.0$ | 2899 | 85 | 10 | 2994 |
| $\Delta\Delta G_{exp} < 2.0$ | 329 | 43 | 5 | 377 |
| $\Delta\Delta G_{exp} > 2.0$ | 71 | 22 | 2 | 95 |
| **total** | 3299 | 150 | 17 | |

Table S2: Performance metrics for smina. Shown are the Pearson correlation coefficient (PCC; higher is better), the mean unsigned error (MUE; lower is better), the root mean squared error (RMSE; lower is better), and the area under the receiver operating characteristics (AUROC; higher is better). Note that smina sometimes produced positive scores. Hence, the MUE and RMSE can get very high for some targets.

| Protein name | PCC | MUE [kcal/mol] | RMSE [kcal/mol] | AUROC |
|---|---|---|---|---|
| Tyrosine-protein kinase JAK2 | 0.43 | 1.89 | 2.15 | 0.72 |
| Estrogen receptor alpha | 0.46 | 1.59 | 2.37 | 0.75 |
| Glucocorticoid receptor | 0.16 | 1.57 | 1.93 | 0.64 |
| Beta-2 adrenergic receptor | 0.25 | 1.69 | 2.05 | 0.60 |
| Androgen receptor | 0.17 | 3.87 | 6.01 | 0.66 |
| Dopamine receptor D2 | 0.11 | 1.49 | 1.82 | 0.58 |
| Tyrosine-protein kinase JAK1 | 0.46 | 2.12 | 2.31 | 0.75 |
| Substance-P receptor | 0.08 | 1.59 | 1.93 | 0.58 |
| 5HT receptor 1B | 0.43 | 1.34 | 1.68 | 0.70 |
| PPAR$\gamma$ | 0.35 | 1.57 | 2.09 | 0.63 |
| Melatonin receptor 1B | 0.37 | 2.18 | 2.63 | 0.68 |
| Phosphodiesterase 4D | 0.59 | 1.47 | 1.96 | 0.74 |
| Estrogen receptor beta | 0.25 | 11.12 | 14.70 | 0.69 |
| Phosphodiesterase 10A | 0.22 | 2.14 | 2.61 | 0.65 |
| Mean | $0.31 \pm 0.15$ | $2.48 \pm 2.59$ | $3.09 \pm 2.70$ | $0.67 \pm 0.06$ |

Table S3: Performance metrics for Glide. Shown are the Pearson correlation coefficient (PCC; higher is better), the mean unsigned error (MUE; lower is better), the root mean squared error (RMSE; lower is better), and the area under the receiver operating characteristics (AUROC; higher is better). Note that Glide failed to dock certain compounds. In these cases, the docking score was set to 0.

| Protein name | PCC | MUE [kcal/mol] | RMSE [kcal/mol] | AUROC |
|---|---|---|---|---|
| Tyrosine-protein kinase JAK2 | 0.06 | 3.32 | 3.73 | 0.53 |
| Estrogen receptor alpha | 0.56 | 1.85 | 2.54 | 0.73 |
| Glucocorticoid receptor | 0.14 | 1.87 | 2.45 | 0.57 |
| Beta-2 adrenergic receptor | 0.37 | 1.57 | 1.97 | 0.65 |
| Androgen receptor | 0.27 | 3.68 | 5.18 | 0.69 |
| Dopamine receptor D2 | 0.09 | 1.60 | 2.01 | 0.60 |
| Tyrosine-protein kinase JAK1 | -0.03 | 3.72 | 4.16 | 0.47 |
| Substance-P receptor | 0.19 | 2.10 | 2.63 | 0.67 |
| 5HT receptor 1B | 0.51 | 1.59 | 1.93 | 0.71 |
| PPAR$\gamma$ | 0.13 | 2.89 | 3.64 | 0.58 |
| Melatonin receptor 1B | 0.34 | 2.25 | 2.86 | 0.69 |
| Phosphodiesterase 4D | 0.02 | 2.31 | 2.60 | 0.52 |
| Estrogen receptor beta | 0.31 | 7.31 | 8.26 | 0.69 |
| Phosphodiesterase 10A | 0.29 | 2.55 | 3.02 | 0.77 |
| Mean | $0.23 \pm 0.18$ | $2.76 \pm 1.51$ | $3.36 \pm 1.69$ | $0.63 \pm 0.09$ |

Table S4: Performance metrics for LeDock. Shown are the Pearson correlation coefficient (PCC; higher is better), the mean unsigned error (MUE; lower is better), the root mean squared error (RMSE; lower is better), and the area under the receiver operating characteristics (AUROC; higher is better).

| Protein name | PCC | MUE [kcal/mol] | RMSE [kcal/mol] | AUROC |
|---|---|---|---|---|
| Tyrosine-protein kinase JAK2 | 0.49 | 2.87 | 3.10 | 0.79 |
| Estrogen receptor alpha | 0.53 | 2.54 | 2.96 | 0.74 |
| Glucocorticoid receptor | 0.31 | 3.24 | 3.56 | 0.69 |
| Beta-2 adrenergic receptor | 0.29 | 2.01 | 2.46 | 0.59 |
| Androgen receptor | 0.41 | 3.48 | 3.84 | 0.71 |
| Dopamine receptor D2 | 0.20 | 2.25 | 2.61 | 0.63 |
| Tyrosine-protein kinase JAK1 | 0.62 | 2.74 | 2.89 | 0.81 |
| Substance-P receptor | 0.01 | 3.41 | 3.88 | 0.63 |
| 5HT receptor 1B | 0.47 | 2.66 | 3.03 | 0.70 |
| PPAR$\gamma$ | 0.40 | 1.65 | 2.04 | 0.69 |
| Melatonin receptor 1B | 0.17 | 4.76 | 5.11 | 0.57 |
| Phosphodiesterase 4D | 0.40 | 2.34 | 2.82 | 0.70 |
| Estrogen receptor beta | 0.20 | 5.36 | 5.77 | 0.62 |
| Phosphodiesterase 10A | 0.59 | 4.20 | 4.49 | 0.82 |
| Mean | $0.36 \pm 0.18$ | $3.11 \pm 1.06$ | $3.47 \pm 1.06$ | $0.69 \pm 0.08$ |

Table S5: Performance metrics for gnina. Shown are the Pearson correlation coefficient (PCC; higher is better), the mean unsigned error (MUE; lower is better), the root mean squared error (RMSE; lower is better), and the area under the receiver operating characteristics (AUROC; higher is better).

| Protein name | PCC | MUE [kcal/mol] | RMSE [kcal/mol] | AUROC |
|---|---|---|---|---|
| Tyrosine-protein kinase JAK2 | 0.65 | 1.53 | 1.77 | 0.87 |
| Estrogen receptor alpha | 0.49 | 2.07 | 2.60 | 0.71 |
| Glucocorticoid receptor | 0.33 | 1.23 | 1.48 | 0.70 |
| Beta-2 adrenergic receptor | 0.32 | 1.54 | 1.86 | 0.63 |
| Androgen receptor | 0.49 | 1.57 | 1.92 | 0.57 |
| Dopamine receptor D2 | 0.23 | 1.13 | 1.39 | 0.61 |
| Tyrosine-protein kinase JAK1 | 0.39 | 1.55 | 1.72 | 0.73 |
| Substance-P receptor | 0.08 | 1.54 | 1.86 | 0.67 |
| 5HT receptor 1B | 0.44 | 1.25 | 1.48 | 0.71 |
| PPAR$\gamma$ | 0.40 | 1.65 | 2.04 | 0.69 |
| Melatonin receptor 1B | 0.22 | 2.32 | 2.78 | 0.62 |
| Phosphodiesterase 4D | 0.50 | 2.09 | 2.52 | 0.68 |
| Estrogen receptor beta | 0.06 | 2.08 | 2.47 | 0.50 |
| Phosphodiesterase 10A | 0.53 | 1.56 | 1.94 | 0.75 |
| Mean | $0.37 \pm 0.17$ | $1.65 \pm 0.36$ | $1.99 \pm 0.44$ | $0.67 \pm 0.09$ |

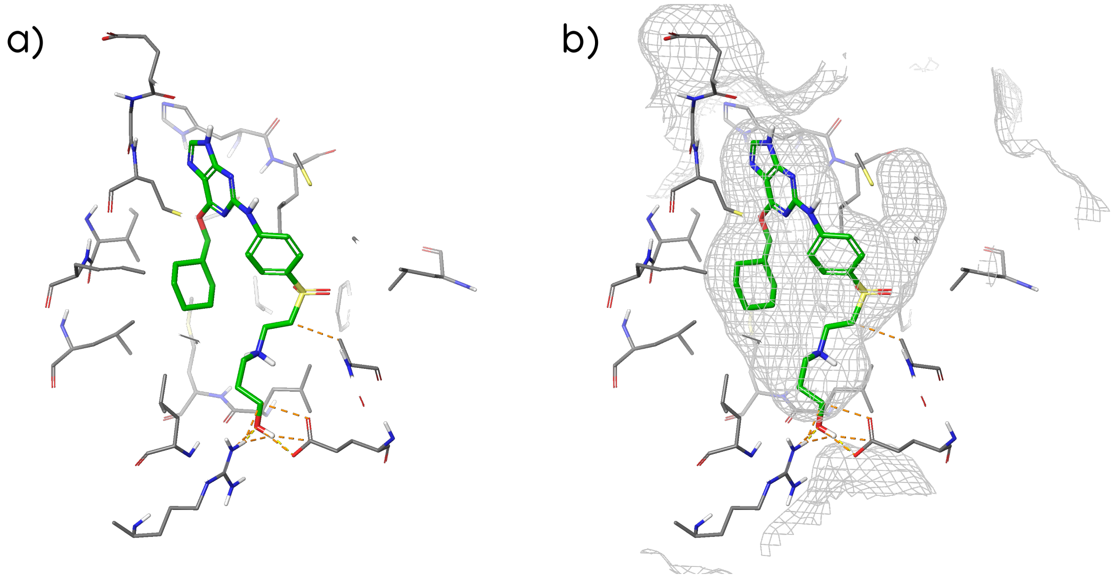


Figure S1: Binding mode of DB08309 at the estrogen receptor alpha generated by Glide. a) without and b) with the binding site surface displayed as a mesh.

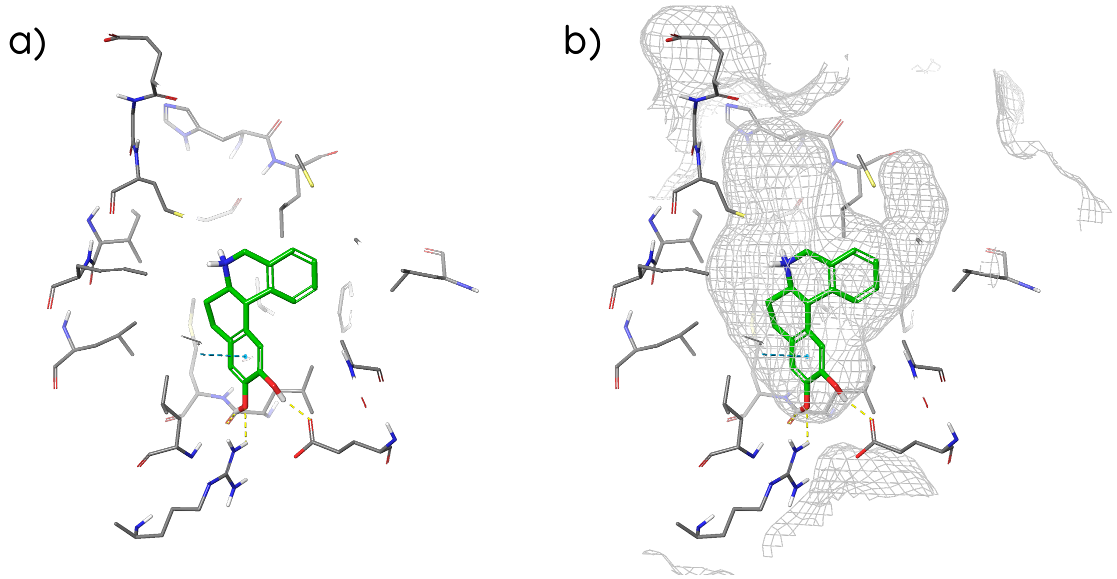


Figure S2: Binding mode of DB16139 at the estrogen receptor alpha generated by Glide. a) without and b) with the binding site surface displayed as a mesh.

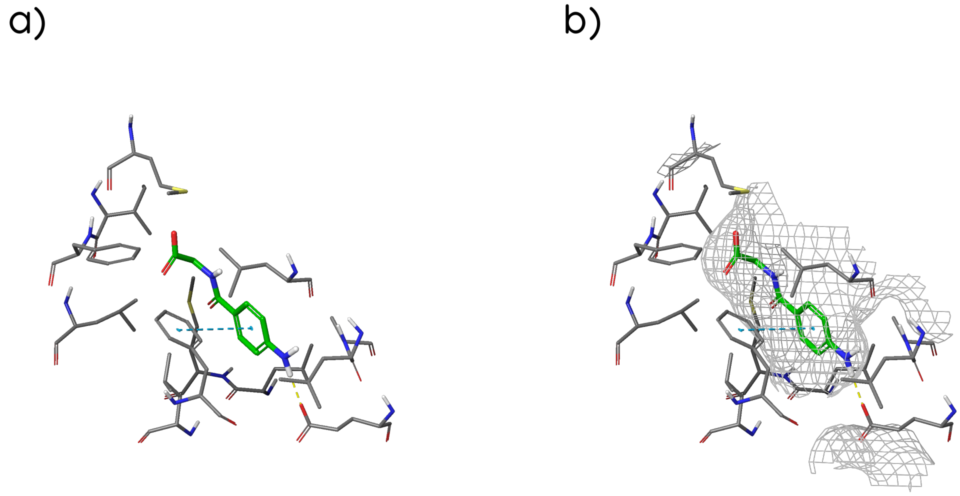


Figure S3: Binding mode of DB00345 at the estrogen receptor alpha generated by Glide. a) without and b) with the binding site surface displayed as a mesh.

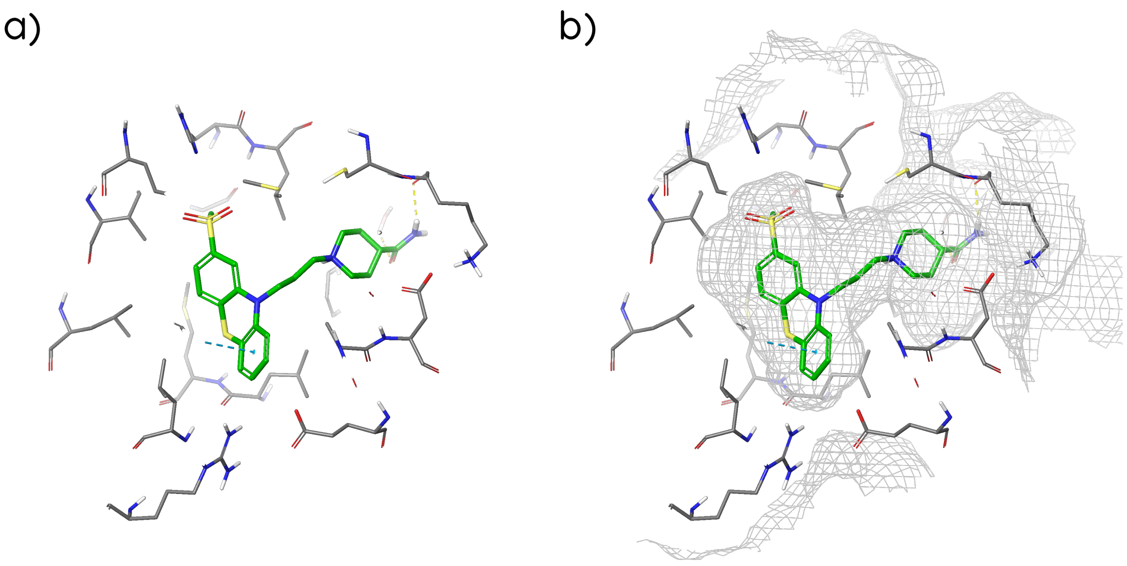


Figure S4: Binding mode of DB13591 at the estrogen receptor alpha generated by Glide. a) without and b) with the binding site surface displayed as a mesh.

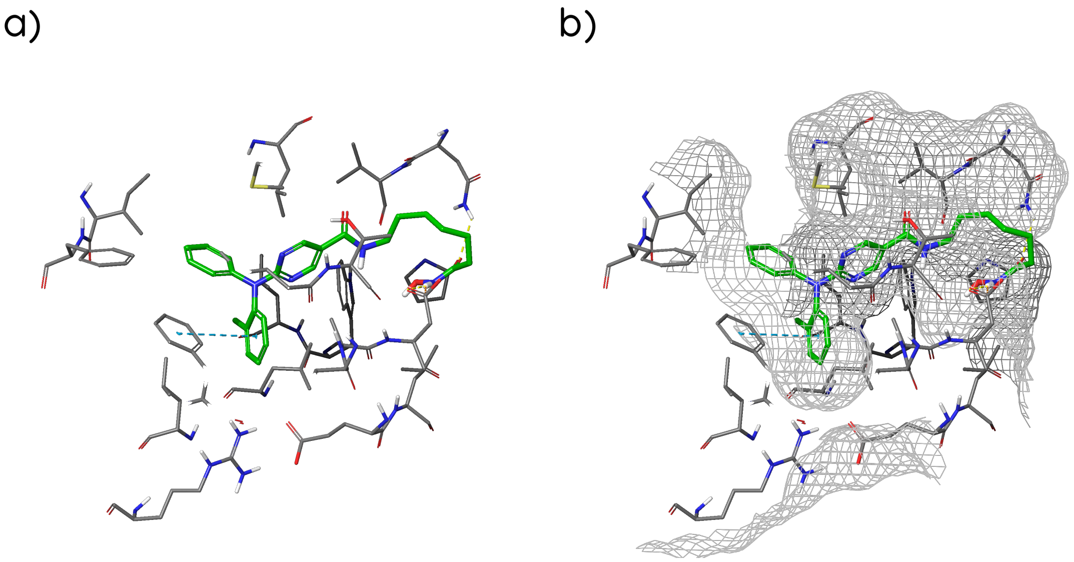


Figure S5: Binding mode of DB15449 at the estrogen receptor alpha generated by Glide. a) without and b) with the binding site surface displayed as a mesh.

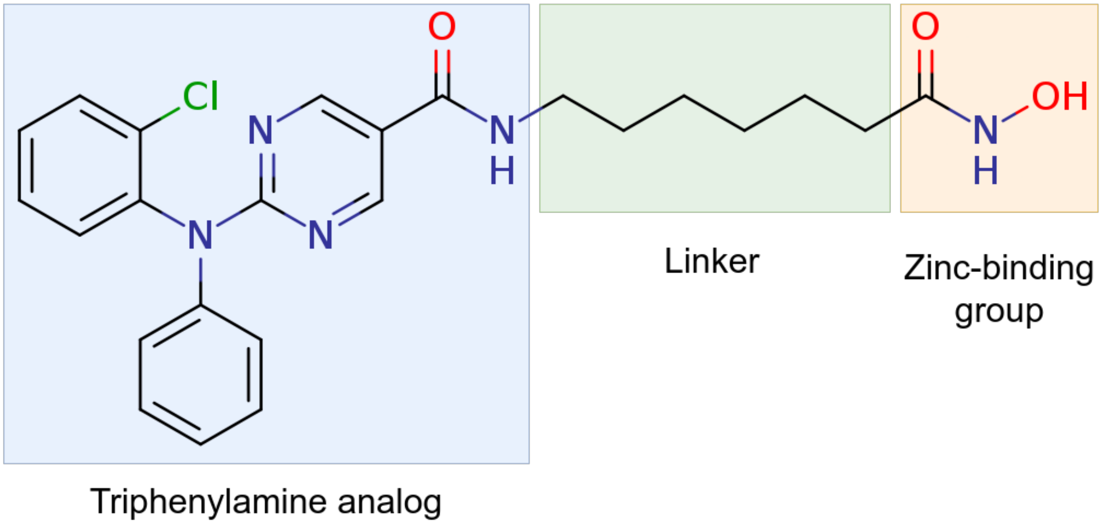


Figure S6: 2D structure of citarinostat.

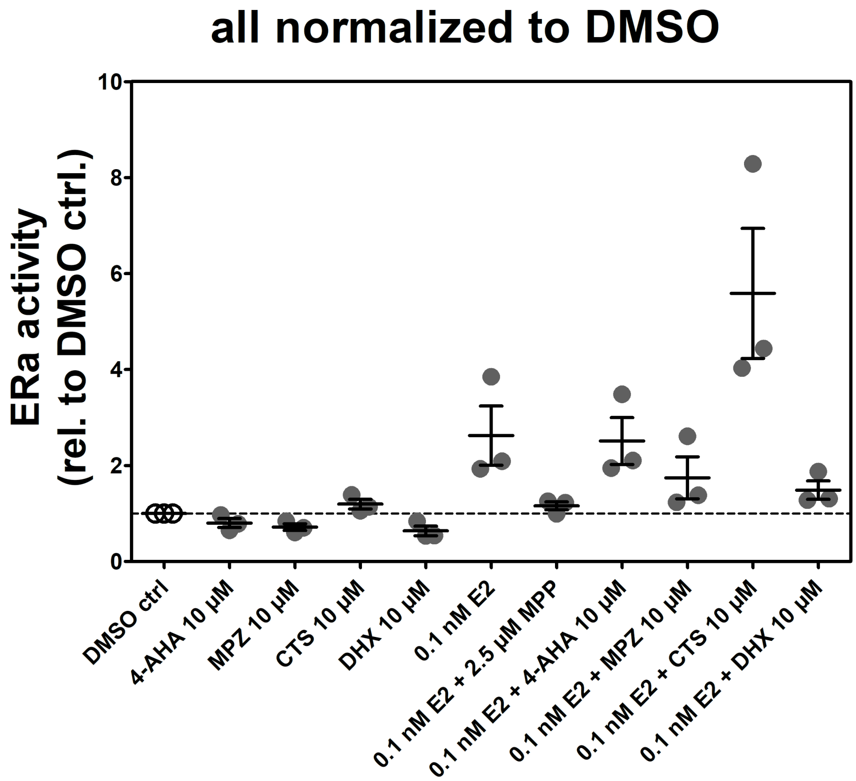


Figure S7: Summary of estrogen receptor alpha activity assay (cell-based). E2: 17$\beta$-estradiol; DMSO: dimethyl sulfoxide; MPP: methyl-piperidino-pyrazole (positive control with known estrogen receptor-alpha antagonsim); 4-AHA: 4-Aminohippuric acid; MPZ: Metopimazine; CTS: Citarinostat; DHX: Dihydrexidine.

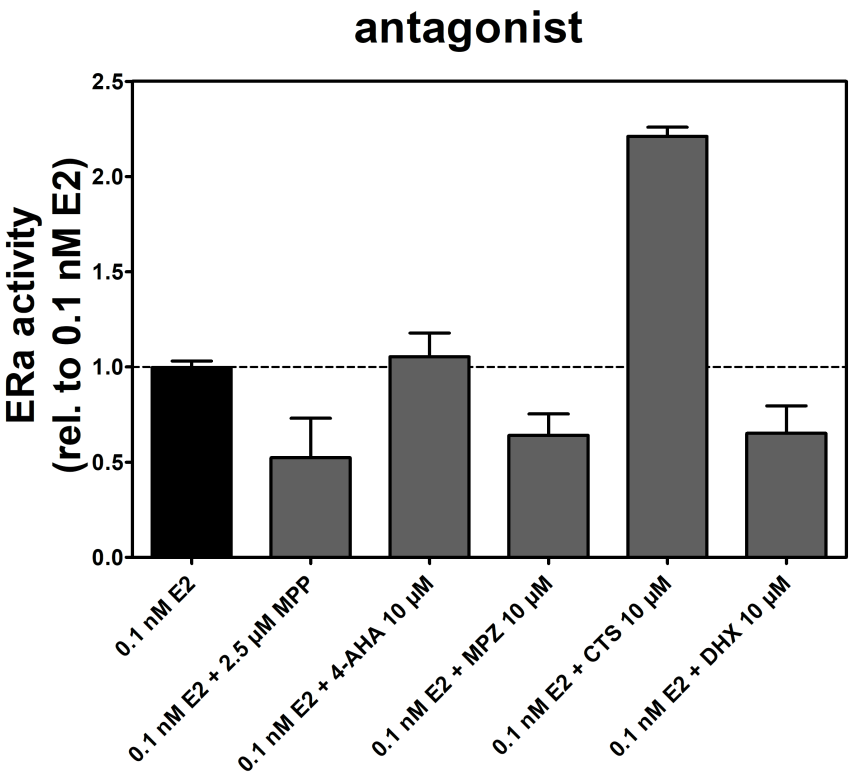


Figure S8: Results of the antagonist-specific estrogen receptor-alpha assay. E2: 17$\beta$-estradiol; MPP: methyl-piperidino-pyrazole (positive control with known estrogen receptor-alpha antagonsim); 4-AHA: 4-Aminohippuric acid; MPZ: Metopimazine; CTS: Citarinostat; DHX: Dihydrexidine.

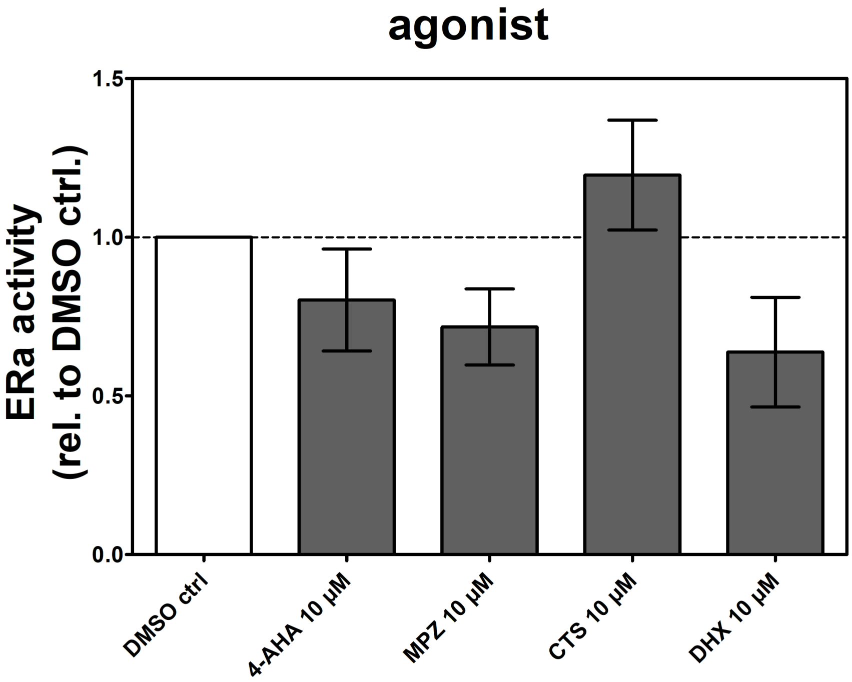


Figure S9: Results of the agonist-specific estrogen receptor-alpha assay. DMSO: dimethyl sulfoxide; 4-AHA: 4-Aminohippuric acid; MPZ: Metopimazine; CTS: Citarinostat; DHX: Dihydrexidine.
